# Expect to neglect: Cross-modal resource allocation in anticipation of visual load

**DOI:** 10.1101/186411

**Authors:** Katrin A. Bangel, Heleen A. Slagter, Ali Mazaheri

## Abstract

Human information processing is limited in capacity. To prevent sensory overload, expectation of upcoming events has been suggested to allocate processing resources to task-relevant regions (e.g., visual system), at the expense of processing in task-irrelevant regions (e.g., auditory system). In support of this, for tasks involving a high visual perceptual load (e.g. visual target search within physically similar distractors), auditory evoked responses were found to be attenuated^1^. This EEG study aimed to further elucidate the neural mechanisms by which the brain prepares for sensory overload. We investigated how expectancy about visual load modulated neural activity, prior to the onset of visual stimuli. Visual load in a letter search task was manipulated by varying the target letter’s similarity to the remaining letters and the letter set size from which flankers were randomly drawn. Importantly, audio-visual cues signaled the likely visual load of the upcoming stimulus-array, manipulating expectancy about visual task load. Cues signaling high visual load elicited attenuated auditory-evoked responses and increased alpha activity over task-irrelevant (auditory) regions, suggesting a functional inhibition of those regions already prior to the arrival of the visual array to suppress auditory cue processing. We also observed a sustained posterior positivity in the ERPs after high perceptual load cues, whose amplitude correlated with reaction times, suggestive of resource allocation for the upcoming visual targets. Expectation about visual load may thus prepare the attentional system both by facilitating target processing and task execution and inhibiting irrelevant sensory processing, thus providing efficient means to overcome attentional limits in situations with complex visual input.

## 1.0 Introduction

The human brain is limited in its processing capacity and thus, being able to anticipate the complexity of upcoming visual scenes may help determine the allocation of processing resources to facilitate information processing. It is well known that top-down attentional control mechanisms can modulate the impact of sensory input on processing in different neural networks, depending on their relevance for the ongoing task^2^^,^^3^. Here it is widely viewed that during top-down control, processes allocate resources to task-relevant regions while functionally inhibiting task-irrelevant regions. This inhibition of task-irrelevant regions happens at the expense of allocating resources to the task-relevant brain region and is believed to be the underlying the cause for the failure to perceive unattended stimuli during tasks involving a high perceptual load^1^. A recent MEG study showed that during a visual search task, auditory evoked brain responses to brief tones as well as tone detection sensitivity were diminished when tones were presented alongside a high visual perceptual load array, as opposed to low load arrays^1^. Here the processing of the high perceptual load array came at the expense of the auditory cortex. However, it remains to be elucidated whether top-down factors, such as expectation of perceptual load, can already attenuate the task-irrelevant areas prior to the onset of the stimuli. While cueing / manipulation of sensory perceptual load has been found to modulate post-target attentional resource allocation^4^^,^^1^^,^^5^^,^^6^, the pre-stimulus neural allocation of resources in anticipation of visual load has rarely been studied. Here we investigated how expectation about perceptual task load in the visual domain, influenced preparatory activity in task-relevant and irrelevant sensory cortices. We were interested in how neural activity in the auditory domain was modulated by expectancy of visual task load prior to task execution and whether these modulations had any relationship to task performance.

A large body of work shows that attention can facilitate processing of task-relevant inputs by increasing the gain of sensory processing^7^^,^^8^. In the visual spatial domain, this is reflected in larger amplitudes of early evoked responses, particularly the N1 over occipitoparietal channels contralateral to the stimulus being processed^9^^,^^10^. In the auditory domain, changes in neural gain in the auditory cortex usually also show in early amplitude modulations over frontocentral electrodes^11^.

In addition to stimulus evoked activity, changes in alpha-band oscillatory activity in preparation for stimulus processing have also been observed to play a pivotal role in the attentional gating of neural activity^12^^,^^13^. Alpha oscillations have been suggested to act as a filter mechanism. They regulate neural excitability in sensory areas and gate sensory information processing as a function of cognitive relevance^12^^-^^15^. Specifically, a number of studies have reported suppression of alpha activity over brain regions that process task-relevant content, while conversely, alpha power is increased over task-irrelevant regions that might interfere with task performance^13^^,^^15^^-^^21^. Moreover, alpha power can be modulated by temporal and spatial expectation in situations of anticipatory attention^12^^,^^22^. It can be increased when participants listen to stimuli under high cognitive load, i.e. presented against noise^23^. In addition a number of studies have observed suppressed alpha power over visual regions and increase of alpha/ beta activity over auditory regions when subjects attended away from the auditory modality towards the visual domain^12^^,^^24^^,^^25^. These findings support an important role of pre-target alpha activity in sensory gating of information during selective spatial or cross-modal attention.

This study’s objective was to investigate how top-down expectancy about upcoming visual task load influences the timing and strength of neural activity related to task preparation. To this end, participants performed a visual search task in which a non-directional multimodal attentional cue manipulated expectancy about upcoming visual task load^5^^,^^26^, while their brain activity was recorded with EEG. This cue consisted of visual and auditory components. We were interested in expectancy-guided modulations of event-related potentials and oscillatory power in the EEG during cue processing and the consequence of this modulation on the filtering of task-irrelevant information and behavioral performance. Specifically, we hypothesized that after audio-visual cues associated with high visual task load, baseline activity in the visual cortex might be enhanced in order to prepare for execution of the visual task, as reflected by decreased alpha band power over visual scalp regions. Similarly, we expected that activity in the task-irrelevant auditory system would be suppressed, as reflected by an increase alpha power over regions associated with auditory processing. In addition, early cue-related ERPs (i.e. N1/P2 reflecting selective attention to cue) and late ERPs (reflecting task preparation) might vary across cue type. Lastly, we expected these cue-related effects to be related to anticipatory attention and to be beneficial for task performance.

## 2.0 Results

### 2.1 Behavioral Results

#### 2.1.1 Benefits of the perceptual load cue for task performance

Reaction times on congruent trials were significantly faster for validly cued high visual load targets compared to invalidly cued (i.e. low perceptual load cue) high load targets. This was revealed through an interaction between cue load*task load*congruency (*F(1,15)* = 4.75, *p* = .046) and post hoc bonferroni-corrected paired-sample t-tests, *t(*15) = −2.60, corrected-*p* = .020; see Fig. 2 left panel). Accuracy was more strongly affected by target-distractor incongruency after cues signalling a high visual load (vs. low load cues, Fig. 2 right panel). This was independent of visual task load as reflected by an interaction of cue load*congruency, *F*(1,15) = 4.87, *p* = .043 and posthoc bonferroni-corrected paired-sample t-tests that revealed a significant congruency effect for high perceptual load cues, *t(*15) = 4.65, *corrected-p* = 0.00062, as well as for low perceptual load cues, *t(*15) = 3.42, *corrected-p* = .0076. Thus, if the possibility for distraction was minimal due to target-distractor congruency, for high visual task load participants’ response time improved by prior cueing of this high load. In addition, relative to low load cues, after the high perceptual load cues participants’ accuracy was impaired by (incongruent) distractors and improved by congruent distractors. This suggests that participants processed and used those cues to perform the search task. High perceptual load cues speeded up task performance and possibly encouraged participants to adopt task strategies that draw more processing resources to the distractors.

**Figure 1.**
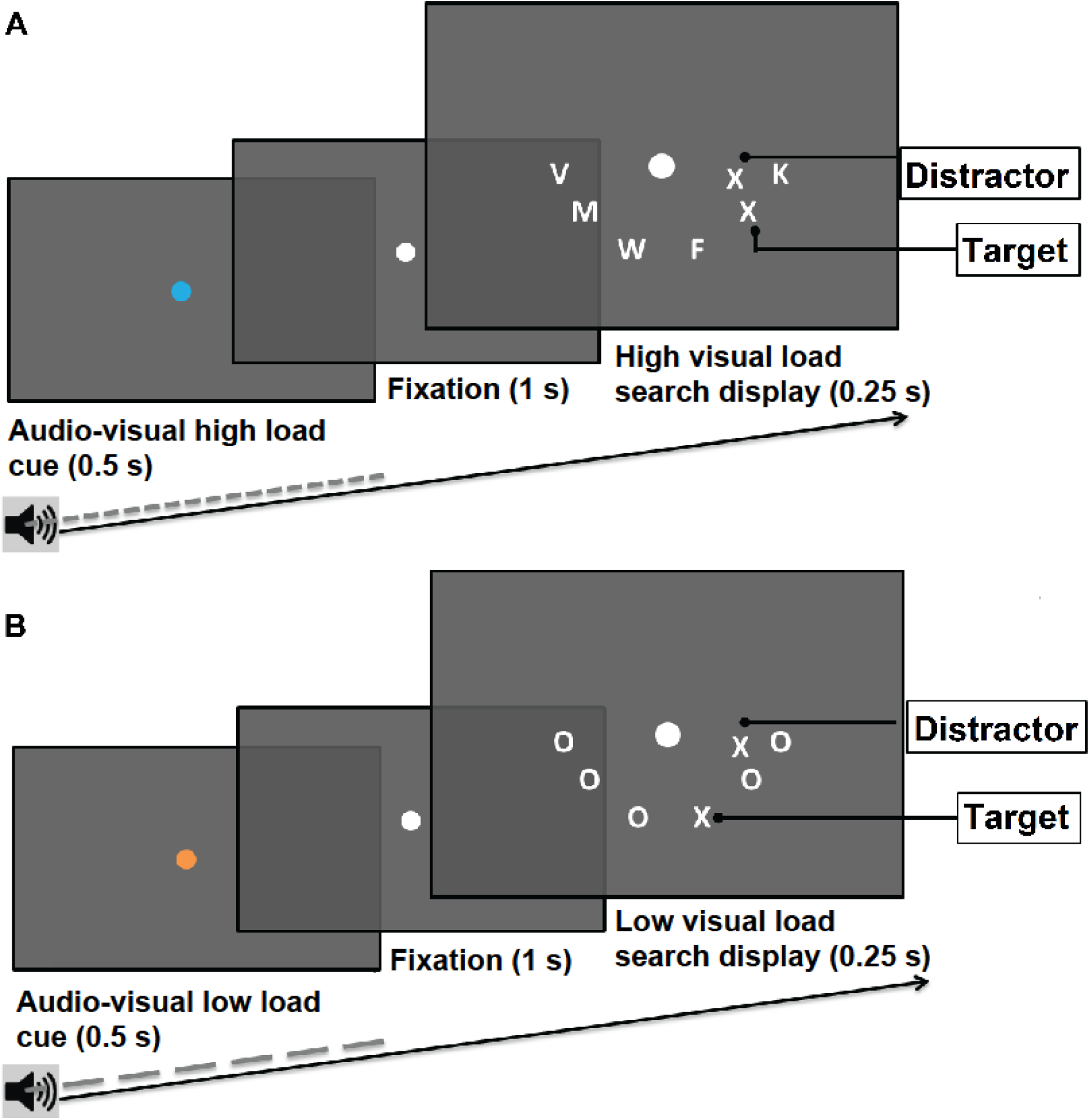
Visual letter search task and examples of possible trial sequences (valid cue & target). Participants discriminated between N or X presented in the arch below fixation. Pre-target audiovisual cues (blue vs. orange, 150 vs. 400 Hz) manipulated expectancy by signalling the likely visual load of the upcoming stimulus-array (85 % validity). Visual task load was manipulated by varying the target letter’s similarity to the remaining letters and by varying the letter set size from which flankers were randomly drawn. Mapping of the cue’s characteristics (color and pitch) to the visual task load (low vs. high) was counterbalanced among participants.

**Figure 2.**
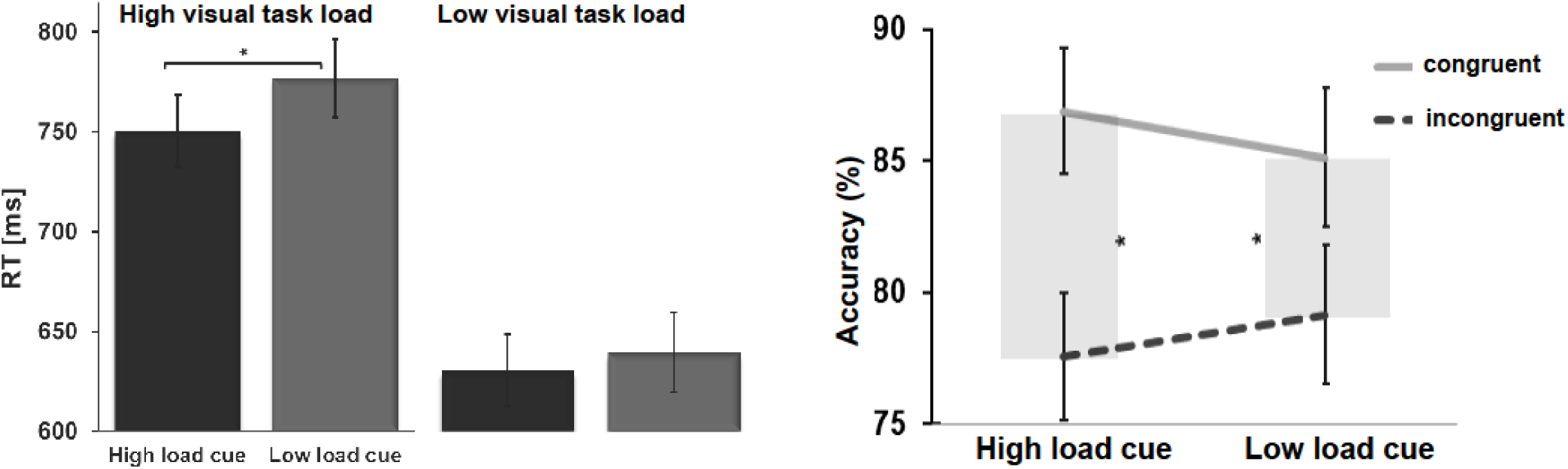
*Left panel*: Reaction times after high/low perceptual load cues and high/low visual task load on congruent trials. Responses to targets with a high visual load were faster after valid high load cues (vs. invalid cues that had signalled a low perceptual load). Here, correctly expecting a high visual load improved task performance, as reflected in faster responses but unimpaired accuracy. *Right panel:* Performance accuracy for high/low perceptual cue loads and target-distractor (in-)congruency. Bold lines illustrate higher accuracy on congruent trials (minimal distraction) compared to incongruent trials with high possibility for distraction (dashed lines), separately for both perceptual load cues. Grey areas reflect the distractor interference on task accuracy that was increased after high vs. low perceptual load cues. Asterisks indicate *p<0.05* (Bonferroni-corrected). The results suggest that participants processed and used the load cues to perform the search task.

#### 2.1.2 Effects of visual task load on task performance

Participants detected low visual task load targets faster and more accurately than high load targets, *F*(1,15) = 63.8, *p* = .0001; low: M = 665.9 ms, SD = 92.5 ms vs. high load: M = 771.2 ms, SD = 103.8 ms and *F*(1,15) = 82.3, *p* < .0001; low: M = 87.9 %, SD = 6.2 % vs high load: M = 76.5 %, SD = 7.4 % respectively. This reaction time and accuracy differences were affected by target-distractor congruency, as reflected by task load*congruency interaction effects on reaction times, *F(1,15)* = 14.75, *p* = .0016 and accuracy, F(1,15) = 6.58, *p* = .0215. A main effect of congruency shows that task performance was generally faster and more accurate when target and distractor letter were congruent, *F*(1,15) = 60.00, *p* = .0001, M = 699.04 ms, SD = 86.01 vs. incongruent: M = 738.19 ms, SD = 95.97 and *F*(1,15) = 19.78, *p* = .0005, M = 86.0 %, SD = 4.5 vs. incongruent: M = 78.4 %, SD = 9.0. Those behavioural effect confirm that our manipulations of visual task load and expectation of load were successful.

### 2.2 EEG Results

#### 2.2.1 Effects of cues signalling likely visual load of stimulus arrays on prestimulus attentional resource allocation: ERPs

We set out to investigate differences in the evoked activity between cues signalling imminent high visual load versus low visual load stimulus arrays. To this end, we examined differences in early cue processing (N1/P2 peak amplitude) and later cue processing in a 1000 ms interval before target onset. The onset of the cue evoked an auditory N1 response at 75 to 125 ms that was maximal at midline central sites, and followed by a late positive complex maximal over posterior cortices. The peak amplitude of the N1 response was more pronounced after low perceptual load cues compared to high perceptual load cues; *t*(13) = 3.15; *p* = .0077 (Fig. 3A). No significant cue evoked differences in P2 were observed. Cluster-based permutation tests revealed two significant differences between high perceptual load versus low perceptual load cues with respect to the late positive complex (Fig. 3). First, from 500 to 565 ms over left centroparietal sensors (Cz, C4, CP1, CP2, Pz, P4, FCz, C1, C2, CPz, CP4, P2, PO4), ERPs showed a stronger positivity for high perceptual load cues, *t*(13) = 39.81; *p* = 0.0001, Monte Carlo *p*-value, corrected for multiple comparisons (Fig. 3B). Second, averages from 590 to 680 ms interval over parietal sensors (P4, P2, P6, P04) were increased after high perceptual load cues, *t*(13) = 13.48; *p* = 0.0001, Monte Carlo *p*-value, corrected for multiple comparisons (Fig. 3C). Post-hoc paired t-tests showed that this latter effect is also significant over a longer interval of 500 to 1250 ms; *t*(13) = 2.17; *p* = .049.

**Figure 3.**
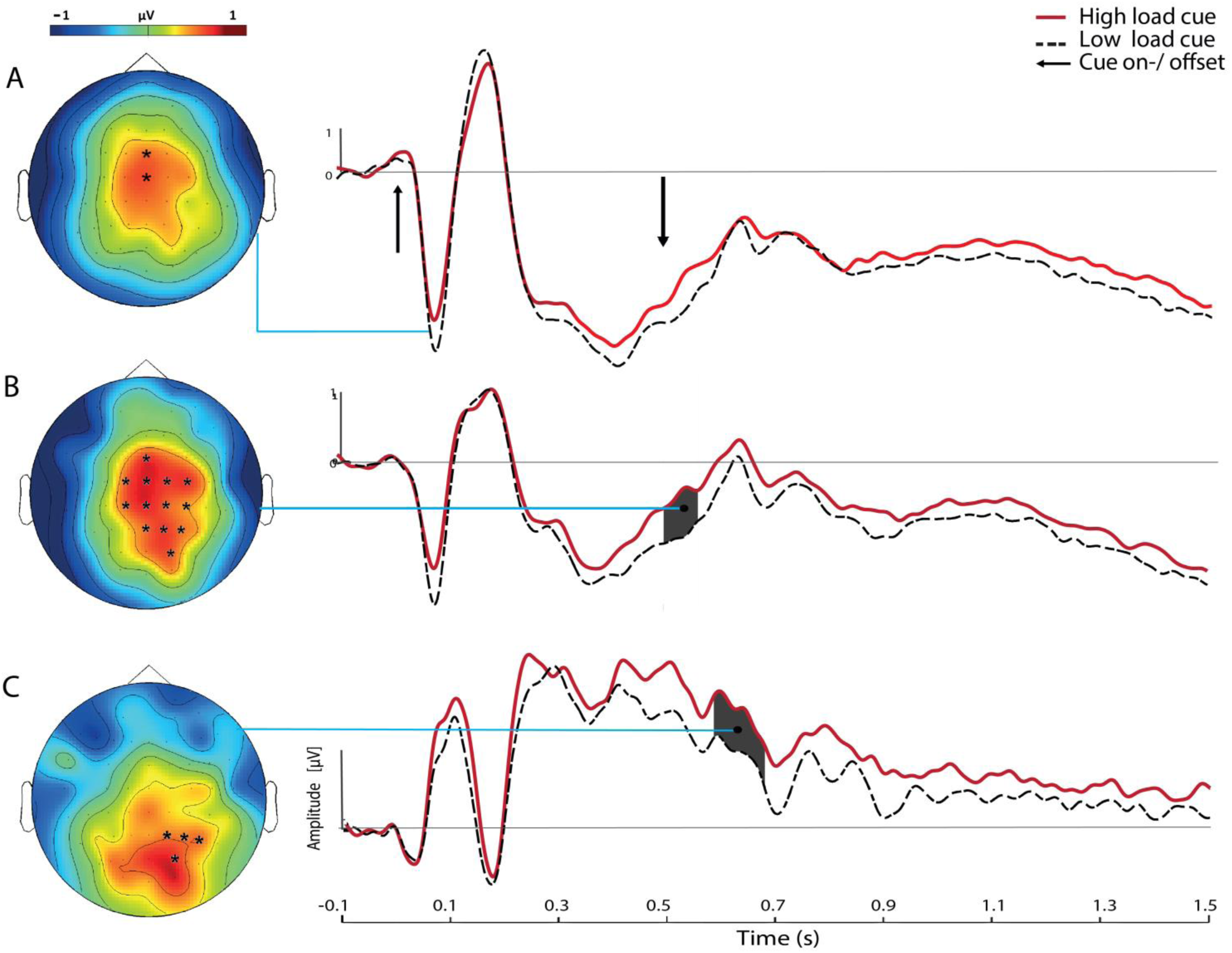
Grand-averaged ERPs contrasting high and low perceptual load cues. (**A)** An attenuated auditory N1 response to high load cues (75 to 125 ms) over a cluster of fronto-central electrodes suggests less pre-attentive processing of the auditory component of high load audiovisual cues. **(B)** Cluster based permutation tests revealed a significantly increased positivity after high load cues over left centroparietal channels (500 to 565 ms post cue, marked in grey). **(C)** This procedure also revealed significantly increased positivity over a cluster of occipito-parietal electrodes (590 to 680 ms post cue, marked in grey). Interestingly, this ERP effect over 590 to 680 ms was significantly related to increased RT benefits from valid high load cues (see Fig. **4**).

#### 2.2.2 Cue-related late posterior positivity correlates with reaction time benefits of the high perceptual load cue

Given that here the late cue-related ERPs were increased for high load cues and the perceptual cue load effect on reaction times was significant in high target congruent trials we performed the following analysis: The late ERPs on high perceptual load cue trials for the two significant clusters were correlated with RT benefit of the perceptual load cue (valid-invalid) for congruent high visual task load trials. ERPs evoked by high perceptual load cues averaged over the second cluster (590 to 680 ms; P4, P2, P6, P04, see Fig. 3C) significantly correlated with RT benefit of the cue (valid-invalid) for high visual task load congruent trials (r_s_= 0.56, df=12, *p* = .037, see Fig. 4), suggesting a direct relation between the cue-related parietal positivity and cue-related reaction time benefits. The correlation of ERPs averaged over the first cluster (Fig 3B) with RT benefit of the cue was not significant.

**Figure 4.**
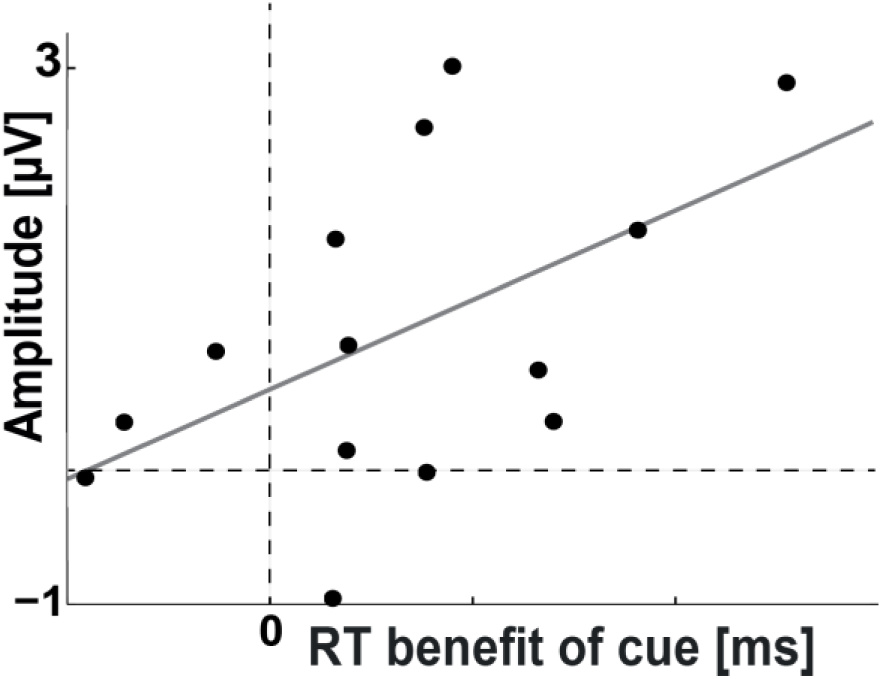
High load cue induced ERP effects of increased late posterior positivity (Fig. **3C**, dark grey) significantly correlated with RT benefits from valid high load cues suggesting enhanced visual attentional capacity during target processing and cue-related behavioral advantages for visual search.

#### 2.2.3 Time-Frequency Analysis: High perceptual load cues induced transient increases in alpha and beta power

Next, we set out to investigate how cues signalling likely high visual load stimulus arrays modulated ongoing alpha activity relative to cues predicting likely low visual load stimulus arrays. Relative to low load cues, high perceptual load cues induced an increase of activity in the alpha band (8-12 Hz, 210-340 ms, *p* =.002, Monte Carlo *p*-value, corrected for multiple comparisons), high alpha/low beta band (12-16 Hz, 210-290 ms, *p* =.002, Monte Carlo *p*-value, corrected for multiple comparisons) and the beta power band (16-20 Hz, 370-410 ms, *p* = .004, Monte Carlo *p*-value, corrected for multiple comparisons) over frontocentral and parietal channels^1^ (see Fig. 5).

**Figure 5.**
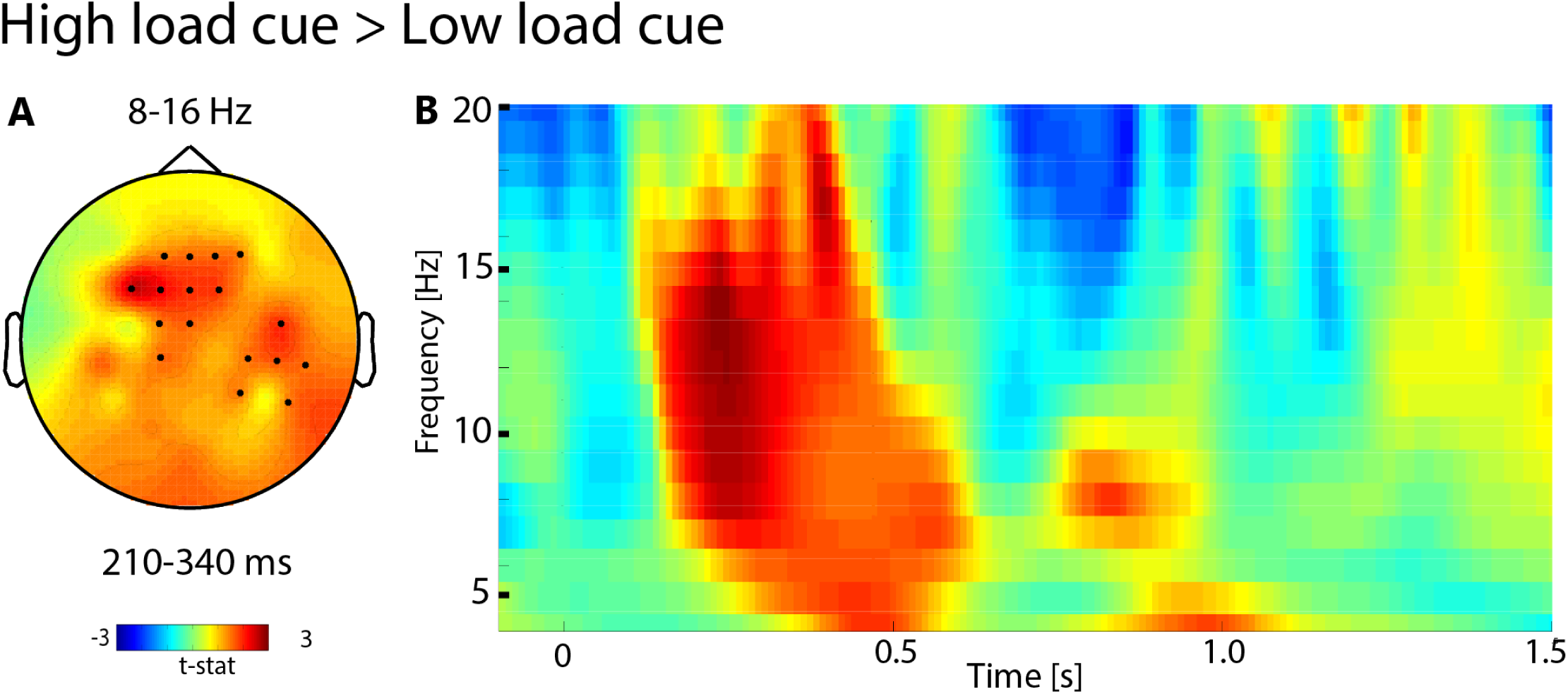
Pre-target perceptual load expectancy modulated alpha/beta power**. (A)** The difference between pre-target power for cues signalling high vs. low visual load revealed a significant increase in alpha (and beta) power over central electrodes in the interval 210-340 ms post cue onset. The differences are expressed in terms of t-values. Electrodes showing the significant modulation (p < 0.05) are marked with black dots. **(B)** Time-frequency representation of oscillatory activity averaged across the significant electrodes.

## 3.0 Discussion

In the current study, we investigated how expectancy about visual load in visual search task modulated the cross-modal allocation of processing resources in the visual and auditory domain. To this end, we utilized a visual search task where visual load was manipulated by varying the target letter’s similarity to the remaining letters and by varying the letter set size from which flankers were randomly drawn. We manipulated expectancy about the visual load through an audio-visual cue, which signalled the likely visual load of the upcoming stimulus-array. Cue effects on behavioral task performance confirmed that our task manipulation induced the intended expectancy effects (Fig. 2). Our findings of faster responses after validly cued high load targets are in line with Spence *et al.* and van Diepen *et al.*^27^^,^^28^. The present findings are the first to illustrate that pre-stimulus task-irrelevant neural activity is supressed prior to stimulus presentation in anticipation of high visual load. We observed that audio-visual cues signalling a high-visual load evoked an attenuated auditory N1 response, suggesting resources were being allocated away from the auditory cortex at a very early stage. Expectancy of high visual load was associated with a sustained posterior positivity in the ERPs (Fig. 3B, 3C) whose amplitude correlated across participants with RT benefits from valid high perceptual load cues (Fig. 4). This sustained positivity may reflect increased gain of neurons in visual brain regions as a response to (visual) cue evaluation. Such cue-evoked gain increases might boost processing capacity for the upcoming more complex, high-load visual task stimuli. Finally, we observed that cues signalling a high-visual load induced an increase in pre-stimulus alpha activity (8-16 Hz) over channels where the N1 was the most prominent. These patterns suggest a disengagement of the auditory processing stream. Expectancy about perceptual load may thus have enhanced visual attentional capacity during target processing, perhaps in part through suppression of irrelevant (auditory) processing.

The observed sustained positivity began over frontocentral channels (Fig. 3B), and over time travelled towards occipital regions, being most pronounced over the (right) visual cortex (Fig. 3C). Neuroimaging, neural stimulation, and single-unit recording studies suggest that a fronto-parietal network plays an essential role in top-down control of attention^29^^-^^34^ and directing attention to relevant stimulus features in preparation of upcoming target information^35^^,^^36^. Specifically, research has shown that the parietal cortex is engaged in shifts of visual attention, and visual stimulus features are encoded via projections to posterior occipital regions^30^^,^^37^. The sustained increases of posterior positivity right after offset of the cue signalling high audio-visual load (Fig. 3C) might thus reflect switches of attention - possibly across sensory modalities - in the course of adapting the attentional system to the expected high visual load of upcoming targets. This notion is supported by the observed relation of this ERP effect with faster response times. Thus far, cue-locked sustained posterior positivity has been found after target-specific attentional cues (e.g., directing attention to color) that reflect cue interpretation^38^^-^^40^, after cues related to task switch/ task set preparation^41^^-^^45^, and lateralized posterior positivity after directional cues related to visual spatial attention / expectation (e.g. LDAP by Hopf & Mangun^46^). Those positive ERPs are typically preceded by negative peaks reflecting location - or feature specific attentional facilitation (e.g. EDAN or switch negativity^7^^,^^39^^,^^46^^-^^49^). Here, expectancy about task load was selectively associated with a sustained unilateral/central positivity. Since, unlike directional or target specific cues, perceptual load cues do not provide direct constraints on target properties, we propose that the underlying meaning of this ongoing positivity (Fig. 3B,C) could be sustained attention to prepare for upcoming visual input. If a cue predicts a complex visual scene, but does not constrain target location or target properties, resources may be allocated for an overall facilitation of visual information processing. Hence, the cueing of upcoming task load might help to preserve our attention in the relevant modality and domain (e.g., visual vs. auditory; global pattern vs. local target attributes).

Task switching studies suggest that the late cue-locked posterior positivity evoked by a task switch cue (e.g., classifying letters instead of numbers) vs. a task repeat cue point towards a dissociation between switch-related and general task preparation subcomponents, with the latter being related to task set reconfiguration^41^^-^^45^. Similar to switching between cognitive tasks, in our experiment preparing for a higher task load might require additional cognitive processing for task set (Fig. 2). Interestingly, a recent pattern classification study associated pre-target right parietal alpha power activation with task readiness / task preparation, but not with task switching^50^. At a longer latency the sustained positivity observed here was more prevalent over the right (visual) cortex. This could hence suggest that expectancy of high load also induced greater general task readiness, but alternatively could also be explained by dominance of the right hemispheric for directing of visual attention^51^^,^^52^ and pre-target attentional biasing^39^^,^^48^.

Expectancy of perceptual load was furthermore associated with an increase in alpha/ beta activity (starting around 210 / 370 ms) over areas that are typically active during selective auditory attention^11^ (see Fig. 5). Pre-target alpha and beta are known to play a role in top-down sensory gating between modalities^25^^,^^53^^-^^55^. Therefore, the observed alpha/beta band synchronisation most likely indexes preparatory blocking of auditory processing. This blocking effect in anticipation of high visual load might serve as a means to allocate cognitive resources to the task-relevant visual regions.

It has been shown that alpha suppression can be invoked cross-modally by auditory symbolic cues^24^. Modality-specific attention can limit processing of stimuli in the unattended sensory modality^56^^,^^57^. Directing attention to the visual domain can attenuate auditory processing^58^^,^^59^, and vice versa^24^. In the present study, we observed blocking of task-irrelevant activity if a high visual load was expected. We speculate that while information from both cue modalities might be integrated in anticipation of low visual load, if a high visual task load is expected, task execution draws upon visual cue information and the auditory task-irrelevant modality is suppressed. As a consequence already before task onset more processing resources can be allocated to the task-specific and relevant modality. This interpretation is supported by our ERP results of supressed auditory N1 after high load signalling cues (Fig 3)^60^^-^^62^. It is also in line with findings of enhanced alpha power during auditory input (speech) under high cognitive load (noise)^23^. Altogether these findings support the notion that alpha synchronization reflects active attentional suppression, and can act as a supramodal mechanism for gating auditory and visual information processing.

Interestingly, we did not observe enhanced alpha suppression over visual areas in preparation for a high visual load. In accordance with findings of Sy *et al.*^5^, if a low task load was expected, attentional capacity might have been used for deeper cue processing (and possible integration of modalities). Yet this is not what was observed and we cannot draw any conclusions based on our data.

In summary, anticipation of task load can prepare the attentional system and thereby modulate resource allocation and task execution. For complex visual tasks, non-spatial warning cues that announce high upcoming visual load may also guide attention by attenuating neuronal processing in task-irrelevant auditory areas and thereby facilitate visual stimulus processing. Providing aids to prepare for the difficulty of an upcoming task thus seems an efficient means to overcome neural processing limits in situations with complex visual input.

## 4.0 Methods

### 4.1 Participants

18 right-handed volunteers were paid 10 €/h for their participation (6 male, mean age = 23.1, age range: 18-27). Participants reported no neurological impairments or any other psychiatric disorders and normal or corrected to normal vision. All participants signed informed consent documents before the experiment. The experiment conformed with World Medical Association Declaration of Helsinki, and was approved by the research ethics committee of the University of Amsterdam.

### 4.2 Design and Procedure

#### 4.2.1 Apparatus and Stimuli

Stimulus presentation was controlled by Presentation software (Neurobehavioral Systems Inc., Albany, CA, USA). A white fixation circle was presented 0.47° below the centre (0.95° in diameter). A perceptual load cue predicted upcoming visual task load and consisted of an auditory cue (150 or 400 Hz) presented together with a visual cue (the fixation circle changing color to blue (0,104,245) or orange (255,130,0)). A search display contained six letters - one target (X or N) and five flankers - which were all white upper case letters (0.95°) presented against a black background in the lower visual field on an arch 3.34° from the center (3.34°, 3.10°, and 2.86° from fixation and 1.91° from each other). Flanker identity was randomly selected from a set of letters with set size depending on visual task load. The search display also contained one distracting letter (white, 0.71°) which was presented in the lower visual field 2.39° to the left or right of fixation (with 0.95° distance from horizontal meridian).

We utilized a paradigm similar to the visual search tasks applied in Lavie and Cox ^63^ and Sy *et al.*^5^. The search task’s load was manipulated by varying the target letter’s similarity to the remaining letters and by varying the letter set size from which flankers were randomly drawn. Low load targets contained five flankers with the same identity. Identity was determined by randomly drawing from a set of four letters (CCCCC, GGGGG, OOOOO or QQQQQ) whose features highly differed from the potential targets (X or N). Flankers in high visual load trials had similar features as the target letter (F, H, K, L, M, T, V, W, Y, Z) and all had unique identities (e.g. YMFZK). At the start of each trial an audio-visual cue predicted the upcoming task load with 85 % validity. The mapping of the cue’s characteristics (color and tone pitch) was counterbalanced to the condition’s visual task load (low vs. high load) across subjects. Third, one to-be-ignored letter was presented unilaterally as a distractor either to the fixation circle’s left or right side. Distractors were presented to the left or right of the fixation circle with equal probabilities and were either congruent (e.g. N target, N flanker) or incongruent (e.g. N target, X flanker) with the presented target’s identity. Congruency of distractor/ flanker presentation side and visual task load (high vs. low visual load) were counterbalanced. Only the likelihood of visual task load, but not likelihood of distractor congruency, were predicted by the perceptual load cues. All other factors were randomly intermixed within each block. An example sequence of a high load and a low load trial is illustrated in Fig. 1.

#### 4.2.2 Experimental Procedure

Each trial started with the load cue for 500 ms, which predicted the visual task load of the upcoming search display. Then the white fixation circle appeared on the screen for 1000 ms. It was followed by a high or low load search display consisting of six letters presented in an arch below the fixation circle for 250 ms. Only the white fixation circle was shown for another 1250 ms while participants responded to the stimulus by button press. To indicate the end of this response interval (of 1500 ms) an exclamation mark was presented at center at the end of each trial. Each participant performed eight blocks of 120 trials each. Participants were asked to report whether the target letter was an X or N, to ignore the distracting letter and to maintain fixation throughout the trial. We also emphasized to make active use of the information provided by the cue on each trial.

### 4.3 Electrophysiological Recordings

EEG was recorded using a WaveGuard 10-20 cap system developed by ANT, with 64 shielded Ag/AgCl electrodes (Advanced Neuro Technology B.V., Enschede, NL). To record horizontal and vertical EOG, electrodes were applied to the outer canthi of the eyes and between supraorbital and infraorbital around the right eye. EEG was continuously sampled at 512 Hz with an online average reference. All recordings were done with ASA software (Advanced Neuro Technology B.V., Enschede, NL).

### 4.4 Data Analysis: Effects of Cue Load

Two participants’ accuracy rates were below 55 % and their data were excluded from the analyses. The EEG data of two additional participants were excluded from the EEG analyses due to extensive movement artifacts. Trials with correct responses to targets were assigned to different conditions based on perceptual cue load (i.e., high vs. low), visual task load (high vs. low), and target-distractor congruency (congruent vs. incongruent).

#### 4.4.1 Behavioral Analysis

For all conditions, accuracy rates [accuracy = number of correct trials/ (number of correct + incorrect trials)] and reaction time of correct trials were calculated. Two repeated-measures (RM)ANOVAs were computed in IBM SPSS (v.20.0) to test the effects of cue load, task load, and flanker-target congruency on reaction time, and accuracy.

#### 4.4.2 EEG Data Analysis

To remove artifactual activity, EEG signal pre-processing was performed using EEGlab 13.3.2b^64^. The data were band-pass filtered between 0.1 and 40 Hz, epoched into 4000 ms windows, and time-locked to the search target stimulus (–2000 to 2000 ms). Subsequently, epochs were checked for large artifacts. Baseline correction was performed using the 100 ms interval before target presentation. Independent component analysis (ICA) was used to detect and remove ocular artifacts and other biological noise sources. We applied the “runica” algorithm, implemented in EEGlab, using the logistic infomax ICA algorithm^65^ and principal component analysis (PCA) dimension reduction to 30 components. On average 4.14 components were removed. Remaining trials that still contained nonbiological artifacts with amplitudes exceeding ±100 μV, were excluded from the analysis. The mean percentage of rejected trials across subjects was 4.55 %. To investigate the effects of expectancy about visual task load, we focused on contrasting high load cue with low load cue condition.

##### 4.4.2.1 Analysis of Event-Related Potentials

ERPs were computed and baseline corrected based on the interval 100 ms prior to cue onset (-1600 to −1500 ms relative to target onset) for all correct trials for both cue types separately. To determine peak latency for early (N1/P2) effects, fixed time intervals were chosen based on the grand-averaged data. Statistical analysis was limited to channel(s) where the overall mean peak amplitude was maximal. The selection of the electrodes was based on aggregated grand average data collapsed over conditions^66^. To investigate effects of cue load on early sensory processing of cues resulting modulations of ERP components paired-samples *t*-tests on individual mean ERP amplitudes were performed. ERP analysis was performed using the open source FieldTrip toolbox for Matlab^67^ (version 20140306; http://www.ru.nl/fcdonders/fieldtrip).

In addition to the analysis of early cue effects, we also investigated later effects of attentional cue processing. ERPs of the time interval −1000 to 0 ms relative to target onset (equivalent to 500 to 1500 ms after cue onset) were analyzed using a cluster-based randomization test^68^. Effects were verified by t-tests on the data averaged over identified time intervals and clusters and statistics are reported in the results section.

To assess the relation between cue-related amplitude modulations and speed of correct target discrimination, for each participant the Spearman correlation between ERP amplitudes averaged over the location and period of interest and subsequent reaction time benefit of the cue was calculated using MatLab (version 2013b, The MathWorks Inc, Natick, MA, USA). Statistical significance of correlation coefficients was assessed by t-tests. For all statistical tests we used an alpha level of 5 % as the statistical criterion.

##### 4.4.2.2 Time-Frequency Analysis

A time-frequency decomposition of the EEG data was performed to investigate the temporo-spectral dynamics of oscillatory modulations by cue load. Time-frequency representations (TFR) of power from 2 to 40 Hz in steps of 1 Hz were obtained per trial using a sliding Hanning window of steps of 50 ms with an adaptive size of three cycles over the data (∆T = 3/f), using the Fieldtrip software package^67^. Similar approaches were used by Jensen *et al.*^15^ and Mazaheri *et al.^69^*. Statistical analyses were conducted separately for each of the frequency bands: low theta (3-5 Hz), high theta (5-7 Hz), alpha (8-12 Hz), high alpha/low beta (12-16 Hz), and beta (16-20 Hz). This classifications of frequency bands were based on prior literature as well as looking at the grand average cue-locked time frequency spectra collapsed across conditions^22^^,^^25^^,^^70^^,^^71^. From those frequency bands, power values of the prestimulus time period (-1300 to 0 ms) were subjected to cluster-level randomisation tests^72^ to statistically evaluate the effects of high vs. low perceptual load cue presentation on neural oscillatory activity. This test contrasted both cue conditions for a high number of channel-time pairs; and to control for multiple comparisons the cluster randomization procedure was used^68^. Two-tailed independent *t*-tests were computed for individual channel-time pairs and thresholded at 5 % significance level. Significant pairs were clustered by direction of effect and spatial proximity using the ‘triangulation’-method (which defines neighbouring sensors based on a two-dimensional projection of the sensor position). We computed the sum of all channel-time t-statistics in each cluster. To assess significance at cluster-level the resulting individual cluster statistics were each compared to a randomization null distribution. This was the distribution of the channel - time pair test statistics under the null hypothesis that both conditions remain the same after 1000 grand-average randomizations of the condition-specific averages. We used the proportion from the randomization distribution in which the maximum cluster-level test statistic exceeded the observed cluster-level test statistic (Monte Carlo estimate of the true cluster p-values which control the false alarm rate). Reported results show topographies and statistics after averaging time points containing a cluster of electrodes with *p* < .05 (Monte Carlo corrected for multiple comparisons).

## 5.0 Acknowledgements

A.M. was supported by a Veni grant from The Netherlands Organisation for Scientific Research (NWO), and H.A.S. by an European Research Council grant [ERC-2015-STG_679399]. Thanks to Dirk J.A. Smit for helpful comments on the manuscript.

## 7.0 Author contributions

K.A.B. and A.M. designed experiment, K.A.B. performed research, analysed data and prepared figures and manuscript, A.M. and H.A.S. supervised data analysis and reviewed the manuscript.

## Footnotes

1 (<u>footnote</u>: alpha: Fz, FC1, FC2, Cz, CP1, CP6, P4, P8, F1, F2, FC3, FCz, C1, C2, C6, CPz, CP4, TP8; high alpha/low beta: ‘Fz, F4, FC1. FC2. Cz, T8, F2, FCz. FC4. C6, TP8 and CP5, P7, P3, O1, PO5, PO3, PO7; beta: F3, FC1, FC2, CP6, P4, AF3, F1, F2, FC3, FCz, C1, C2, CP4, P2, P6, PO4, PO6).

## References

[1] Molloy, K., Griffiths, T. D., Chait, M. & Lavie, N. Inattentional Deafness: Visual Load Leads to Time-Specific Suppression of Auditory Evoked Responses. J. Neurosci. 35, 16046–16054 (2015).

[2] Desimone, R. & Duncan, J. Neural mechanisms of selective visual attention. Annu. Rev. Neurosci. 18, 193–222 (1995).

[3] Egeth, H. E. & Yantis, S. Visual attention: Control, representation, and time course. Annu. Rev. Psychol. 48, 269–297 (1997).

[4] Cartwright-Finch, U. & Lavie, N. The role of perceptual load in inattentional blindness. Cognition 102, 321–340 (2007).

[5] Sy, J. L., Guerin, S. A., Stegman, A. & Giesbrecht, B. Accurate expectancies diminish perceptual distraction during visual search. Front. Hum. Neurosci. 8, 334

[6] Otten, L. J., Alain, C. & Picton, T. W. Effects of visual attentional load on auditory processing. Neuroreport 11, 875–880 (2000).

[7] Hillyard, S. a & Anllo-Vento, L. Event-related brain potentials in the study of visual selective attention. Proc. Natl. Acad. Sci. U. S. A. 95, 781–787 (1998).

[8] Luck, S. J., Chelazzi, L., Hillyard, S. a & Desimone, R. Neural mechanisms of spatial selective attention in areas V1, V2, and V4 of macaque visual cortex. J. Neurophysiol. 77, 24–42 (1997).

[9] Mangun, G. R. & Hillyard, S. A. The spatial allocation of visual attention as indexed by event-related brain potentials. Hum. Factors 29, 195–211 (1987).

[10] Slagter, H. A., Prinssen, S., Reteig, L. C. & Mazaheri, A. Facilitation and inhibition in attention: Functional dissociation of pre-stimulus alpha activity. in P1, and N1 components. NeuroImage 125, 25–35

[11] Woldorff, M. G. et al. Modulation of early sensory processing in human auditory cortex during auditory selective attention. in Proceedings of the National Academy of Sciences 90, 8722–8726

[12] Foxe, J. J., Simpson, G. V. & Ahlfors, S. P. Parieto-occipital ~10 Hz activity reflects anticipatory state of visual attention mechanisms. Neuroreport 9, 3929–3933

[13] Klimesch, S. & Hanslmayr. EEG alpha oscillations: the inhibition-timing hypothesis. Brain Res. Rev. 53, 63–88

[14] Jensen, O. & Mazaheri, A. Shaping functional architecture by oscillatory alpha activity: gating by inhibition.

[15] Jensen, O., Gelfand, J., Kounios, J. & Lisman, J. E. Oscillations in the alpha band (9-12 Hz) increase with memory load during retention in a short-term memory task. Cereb. cortex 12, 877–882

[16] Busch, C. S. H. Object-load and feature-load modulate EEG in a short-term memory task. Neuroreport 14, 1721–1724

[17] Cooper, R. J. C., Dominey, S. J., Burgess, A. P. & Gruzelier, J. H. Paradox lost? J. Psychophysiol. 47, 65–74

[18] Herrmann, C. S., Senkowski, D. & Röttger, S. Phase-locking and amplitude modulations of EEG alpha: two measures reflect different cognitive processes in a working memory task. Experimental 51, 311–318

[19] Klimesch, W. EEG alpha and theta oscillations reflect cognitive and memory performance: a review and analysis. Brain Res. Rev 29, 195

[20] S.P, K. Increases in alpha oscillatory power reflect an active retinotopic mechanism for distracter suppression during sustained visuospatial attention. J. Neurophysiol, 95 3844–385

[21] Rihs, T. A., Michel, C. M. & Thut, G. Mechanisms of selective inhibition in visual spatial attention are indexed by α-band EEG synchronization.European. J. Neurosci. 25, 603–610

[22] van Diepen, R. M., Cohen, M. X., Denys, D. & Mazaheri, A. Attention and temporal expectations modulate power, not phase, of ongoing alpha oscillations. J. Cogn. Neurosci.

[23] Wilsch, A., Henry, M. J., Herrmann, B., Maess, B. & Obleser, J. Alpha oscillatory dynamics index temporal expectation benefits in working memory. Cereb. Cortex 25, 1938–1946

[24] Fu, K. M. G. et al. Attention-dependent suppression of distracter visual input can be cross-modally cued as indexed by anticipatory parieto–occipital alpha. Cogn. Brain Res. 12, 145–152

[25] Mazaheri, A. et al. Region-specific modulations in oscillatory alpha activity serve to facilitate processing. in the visual and auditory modalities. Neuroimage 87, 356–362

[26] Lavie, N. Perceptual load as a necessary condition for selective attention. J. Exp. (Psychol.: Human Percept. Perform).

[27] Spence, C., Nicholls, M. E. & Driver, J. The cost of expecting events in the wrong sensory modality. Percept Psychophys 63, 330–336

[28] van Diepen, R. M., Miller, L. M., Mazaheri, A. & Geng, J. J. The Role of Alpha Activity in Spatial and Feature-Based Attention. eneuro 3, 204

[29] Brignani, D., Lepsien, J., Rushworth, M. F. & Nobre, A. C. The timing of neural activity during shifts of spatial attention. J. Cogn. Neurosci. 21, 2369–2383

[30] Corbetta, M., Miezin, F. M., Shulman, G. L. & Petersen, S. E. A PET study of visuospatial attention. J. Neurosci. 13, 1202–1226

[31] Corbetta, M. & Shulman, G. L. Control of goal-directed and stimulus-driven attention in the brain. Nat. Rev. Neurosci. 3, 201–215

[32] Gottlieb, J. From thought to action: The parietal cortex as a bridge between perception, action, and cognition. Neuron 53, 9–16

[33] Hopfinger, J. B., Woldorff, M. G., Fletcher, E. M. & Mangun, G. Dissociating top–down attentional control from selective perception and action. Neuropsychologia 39, 1277–1291

[34] Taylor, P. C., Nobre, A. C. & Rushworth, M. F. FEF TMS affects visual cortical activity. Cereb. Cortex 17, 391–399

[35] Liu, T., Slotnick, S. D., Serences, J. T. & Yantis, S. Cortical mechanisms of feature-based attentional control. Cereb. cortex 13, 1334–1343

[36] Shulman, G. L. et al. Areas involved in encoding and applying directional expectations to moving objects. J. Neurosci. 19, 9480–9496

[37] Desimone, R., Wessinger, M., Thomas, L. & Schneider, W. Attentional control of visual perception: Cortical and subcortical mechanisms. in Cold Spring Harbor Symposia on Quantitative Biology 55, 963–971 (1990).

[38] Slagter, H. A., Kok, A., Mol, N., Talsma, D. & Kenemans, J. L. Generating spatial and nonspatial attentional control: An ERP study. Psychophysiology 42, 428–439

[39] Harter, M. R., Miller, S. M., Price, N. B., LaLonde, M. E. & Keyes, A. L. Neural processes involved in directing attention. Cogn. Neurosci. J. 1, 223–237

[40] Hillyard, S. A. & Munte, T. F. Selective attention to color and location: an analysis with event-related brain potentials. Percept Psychophys 36, 185–198 (1984).

[41] Karayanidis, F., Provost, A., Brown, S., Paton, B. & Heathcote, A. Switch-specific and general preparation map onto different ERP components in a task-switching paradigm. Psychophysiology 48, 559–568 (2011).

[42] Karayanidis, F. et al. Anticipatory reconfiguration elicited by fully and partially informative cues that validly predict a switch in task. Cogn. Affect. Behav. Neurosci. 9, 202–215 (2009).

[43] Karayanidis, F. et al. Advance preparation in task-switching: Converging evidence from behavioral, brain activation, and model-based approaches. Frontiers in Psychology (2010). doi:10.3389/fpsyg.2010.00025

[44] Kieffaber, P. D. & Hetrick, W. P. Event-related potential correlates of task switching and switch costs. Psychophysiology 42, 56–71 (2005).

[45] Lavric, A., Mizon, G. A. & Monsell, S. Neurophysiological signature of effective anticipatory task-set control: A task-switching investigation. Eur. J. Neurosci. 28, 1016–1029 (2008).

[46] Hopf, J.-M. & Mangun, G. Shifting visual attention in space: an electrophysiological analysis using high spatial resolution mapping. Clin. Neurophysiol. 111, 1241–1257 (2000).

[47] Yamaguchi, S., Tsuchiya, H. & Kobayashi, S. Electrophysiologic correlates of age effects on visuospatial attention shift. Cogn. Brain Res. 3, 41–49

[48] Yamaguchi, S., Tsuchiya, H. & Kobayashi, S. Electroencephalographic activity associated with shifts of visuospatial attention. Brain 117, 553–562

[49] Harter, M. R. & Aine, C. J. Brain mechanisms of visual selective attention. in Varieties of attention, pp (eds. Parasuraman, R. & Davies, D. R.) 293–321

[50] Mansfield, E. L., Karayanidis, F. & Cohen, M. X. Switch-Related and General Preparation Processes in Task-Switching: Evidence from Multivariate Pattern Classification of EEG Data. J. Neurosci. 32, 18253–18258 (2012).

[51] Heilman, K. M. & den Abell, T. Right hemispheric dominance for mediating cerebral activation. Neuropsychologia 17, 315–321

[52] Mesulam, M. A cortical network for directed attention and unilateral neglect. Ann. 10, 309–325

[53] Engel, A. K. & Fries, P. Beta-band oscillations—signalling the status quo? in Current opinion in neurobiology 20, 156–165

[54] Min, B. K. & Herrmann, C. S. Prestimulus EEG alpha activity reflects prestimulus top-down processing. Neurosci. Lett. 422, 131–135

[55] van Ede, F., de Lange, F., Jensen, O. & Maris, E. Orienting attention to an upcoming tactile event involves a spatially and temporally specific modulation of sensorimotor alpha-and beta-band oscillations. J. Neurosci. 31, 2016–2024

[56] Mozolic, J. L., Hugenschmidt, C. E., Peiffer, A. M. & Laurienti, P. J. Modality-specific selective attention attenuates multisensory integration. Exp. brain Res. 184, 39–52

[57] Talsma, D., Doty, T. J. & Woldorff, M. G. Selective attention and audiovisual integration: is attending to both modalities a prerequisite for early integration? Cereb Cortex 17, 679–690

[58] Alho, K., Woods, D. L. & Algazi, A. Processing of auditory stimuli during auditory and visual attention as revealed by event-related potentials. Psychophysiology 31, 469–479

[59] Woods, D. L., Alho, K. & Algazi, A. Intermodal selective attention. I. Effects on event-related potentials to lateralized auditory and visual stimuli. Electroencephalogr. Clin. Neurophysiol. 82, 341–355

[60] Luck, S. J. & Hillyard, S. A. Electrophysiological correlates of feature analysis during visual search. Psychophysiology 31, 291–308

[61] Mangun, G. R. Orienting attention in the visual fields: An electrophysiological analysis. in Cognitive electrophysiology 81–101 (Birkhäuser Boston).

[62] Talsma, D., Slagter, H. A., Nieuwenhuis, S., Hage, J. & Kok, A. The orienting of visuospatial attention: An event-related brain potential study. Cogn. Brain Res. 25, 117–129

[63] Lavie, N. & Cox, S. On the efficiency of visual selective attention: Efficient visual search leads to inefficient distractor rejection. Psychol. Sci. 8, 395–396

[64] Delorme, A. et al. EEGLAB, SIFT, NFT, BCILAB, and ERICA: new tools for advanced EEG processing. Comput. Intell. Neurosci. 10

[65] Bell, A. J. & Sejnowski, T. J. An information-maximization approach to blind separation and blind deconvolution. Neural Comput. 7, 1129–1159

[66] Brooks, J. L., Zoumpoulaki, A. & Bowman, H. Data-driven region-of-interest selection without inflating Type I error rate. Psychophysiology 54, 100–113

[67] Oostenveld, R., Fries, P., Maris, E. & Schoffelen, J. M. FieldTrip: open source software for advanced analysis of MEG, EEG, and invasive electrophysiological data. Comput. Intell. 2011,

[68] Maris, E. Randomization tests for ERP topographies and whole spatiotemporal data matrices. Psychophysiology 41, 142–151

[69] Mazaheri, A., DiQuattro, N. E., Bengson, J. & Geng, J. J. Pre-stimulus activity predicts the winner of top-down vs. bottom-up attentional selection. PLoS One 6, 16243

[70] Bengson, J. J., Mangun, G. R. & Mazaheri, A. The neural markers of an imminent failure of response inhibition. Neuroimage 59, 1534–1539

[71] Hanslmayr, S. et al. Visual discrimination performance is related to decreased alpha amplitude but increased phase locking. Neurosci. Lett. 375, 64–68

[72] Maris, E. & Oostenveld, R. Nonparametric statistical testing of EEG-and MEG-data. J. Neurosci. Methods 164, 177–190

